# A spatial atlas of Wnt and Frizzled receptor expression in adult mouse liver

**DOI:** 10.1101/2022.09.23.509269

**Authors:** Jenesis Gayden, Shikai Hu, Paul N. Joseph, Evan Delgado, Silvia Liu, Aaron Bell, Stephanie Puig, Satdarshan P. Monga, Zachary Freyberg

## Abstract

Hepatic zonation is critical for most metabolic functions in liver. Wnt signaling plays an important role in establishing and maintaining liver zonation. Yet, the anatomic expression of Wnt signaling components, including all 10 Frizzled receptors (Fzds), has not been characterized in adult liver. To address this, we quantitatively mapped the spatial expression of Wnt/Fzd pathway components in adult mouse liver via multiplex fluorescent *in situ* hybridization. While all 10 Fzds are expressed within a metabolic unit, Fzds 1, 4, and 6 are the highest expressed. Though the majority of Wnt signaling occurs in zone 3, expression of most Fzds is not zonated. In contrast, Fzd6 is preferentially expressed in zone 1. We also discovered that Wnt2 and Wnt9b expression is highly zonated and primarily found in zone 3. Therefore, our results suggest that zonated Wnt expression is critical for zonation maintenance in healthy adult liver. Finally, we showed that Fzds and Wnts are not uniformly expressed by all hepatic cell types. Rather, there is broad distribution among both hepatocytes and non-parenchymal cells, including endothelial cells. Overall, our establishment of a definitive mRNA expression atlas of Wnt/Fzd pathway components opens the door to future functional characterization in healthy and disease states.

## Introduction

Liver performs several functions critical for metabolic health, including cholesterol metabolism, drug detoxification, and synthetic functions [1, 2]. The ability to compartmentalize these various metabolic functions is an essential aspect of hepatic physiology [3]. Anatomically, this compartmentalization is represented by zonation of the hepatic lobule into distinct zones (zones 1-3) along the periportal-central vein axis. Functionally, zonation guides hepatocytes to metabolic processes specific to each zone (*i*.*e*., “metabolic zonation”) [4-6]. These differences depend on differential expression of enzymes, proteins, transcription factors, and oxygenation which occur along a gradient. For example, periportal hepatocytes in zone 1 predominantly catalyze the oxidative catabolism of fatty and amino acids as well as gluconeogenesis-driven glucose release and glycogen formation [4, 6]. In contrast, pericentral cells in zone 3 preferentially engage in glucose uptake for glycogen synthesis and glycolysis coupled to *de novo* lipogenesis [7]. Significantly, increasing evidence suggests that zone-specific differences in gene expression are essential for establishing and maintaining metabolic zonation [5, 8]. Such zonal division of labor makes liver an efficient metabolic, synthetic, and detoxifying organ.

We and others have established that Wnt signaling is a fundamental regulator of hepatic zonation by controlling zone-specific gene expression [8-11]. Consistent with this, Wnt signaling is zone-dependent and follows the metabolic gradient, activating gene expression in pericentral hepatocytes (zone 3), while suppressing genes in its periportal counterparts (zone 1). Indeed, we showed that Wnt2- and Wnt9b- dependent signaling is essential to maintain zonation [12]. More recently, we conditionally eliminated Wnt2 and Wnt9b expression from hepatic endothelial cells including those lining the central vein and sinusoidal endothelial cells in zone 3 and zone 2. This led to complete disruption of β-catenin activation in these zones and a notable defect in pericentral hepatocyte gene expression [13]. Investigations of the liver’s capacity for injury recovery and regeneration have yielded key insights into the roles of Wnt signaling in metabolic zonation, not only in development, but also in adult tissue homeostasis [6, 8, 14-16]. Previously, we uncovered that 11 Wnts and 8 Frizzled (Fzd) genes were expressed in normal liver [17], suggesting a function for Wnt signaling post-development, including maintenance of metabolic zonation. While much is known regarding hepatic Wnt signaling, the precise anatomic expression of these signaling components remains poorly characterized in adult liver.

Defining the expression patterns of the Wnt signaling machinery is a crucial first step in better understanding the post-developmental roles of a pathway traditionally associated with development. Wnt/β-catenin signaling is largely quiescent in adult liver except in the pericentral region of the hepatic lobule (zone 3). Therefore, we hypothesized that Wnt signaling components are expressed along a gradient with most expression in pericentral zone 3. To test this, we utilized newly developed multiplex RNAscope fluorescent *in situ* hybridization to spatially characterize the mRNA expression of Wnt signaling machinery, including Wnt2, Wnt9b, Frizzled receptors 1-10, and co-receptors low-density lipoprotein receptor-related protein 5 and 6 (Lrp5, Lrp6). Importantly, this enabled us to bypass past limitations in mapping hepatic Wnt signaling at the protein level due to the lack of specific antibodies.

## Methods

### Animals

Animals were housed and handled in accordance with appropriate NIH guidelines through the University of Pittsburgh Institutional Animal Care and Use Committee (protocol# 19126451). We abided by all appropriate animal care guidelines for reporting animal research. Mice were housed in cages with a 12:12 light:dark cycle and had access to food and water *ad lib* at all times.

### Liver preparation

Livers were harvested from male C57BL/6J mice (The Jackson Laboratory, Bar Harbor, ME) between the ages of 2.5-3-months. Liver tissue was washed in ice-cold PBS and processed into chunks no larger than 2mm^2^. Liver chunks were fixed in 4% PFA (overnight, 4°C), incubated in 30% sucrose in PBS (overnight, 4°C) until fully submerged and were OCT-embedded and frozen on dry ice. Tissue blocks were stored at -80°C until used.

### Multiplex fluorescent *in situ* hybridization

*In situ* detection of mRNA expression was attained via fluorescent multiplex RNAscope (Advanced Cell Diagnostics, Hayward, CA) according to manufacturer’s instructions. Fixed frozen mouse liver sections (5µm thick) were first mounted on slides and then submerged in target retrieval solution (5min, 100ºC), followed by protease pretreatment (30min, 40ºC). Slides were then incubated with probes targeting Fzds 1-10, Wnt2, Wnt9b, Lrp5, and Lrp6 (2h, 40ºC). We defined metabolic zones using additional probes glutamine synthetase (GS) and glutaminase2 (Gls2). Slides were then treated with an amplification kit (Advanced Cell Diagnostics) to hybridize each probe to its respective mRNA target, followed by fluorescent labeling (90min, 40ºC). Finally, slides were treated with DAPI (Advanced Cell Diagnostics) to visualize cell nuclei and mounted with Prolong Gold Antifade mountant (Invitrogen, Waltham, MA).

### Confocal microscopy

Images were acquired with an Olympus VS120 whole slide scanner microscope (Olympus America, Center Valley, PA) equipped with an Olympus 20x objective (0.75 NA) and a Hamamatsu ORCA Flash4.0 V2 digital camera (Hamamatsu Corporation, Bridgewater, NJ) (0.33µm/pixel). Image stacks were acquired during scanning.

### Image analysis

Using Olympus VS-ASW software, image stacks were converted into maximum intensity projections and exported as TIFF files. Image analysis was performed using HALO image analysis platform equipped with a fluorescent *in situ* hybridization plug-in (Version 3.0, Indica Labs, Albuquerque, NM). Nuclei were quantified as DAPI-stained objects within the 30-150µm^2^ range and the minimum cytoplasmic radius was set at 5µm. Puncta corresponding to the respective mRNA probes were quantified as any 0.01-0.15µm^2^ object. To account for background, the number of identified puncta for each probe was divided by the total area analyzed. This number was then multiplied by the average cell area, providing an average cellular density per mRNA probe (positive cells/mm^2^). Thus, these thresholds represented 3 times the smaller observed density level and were used as a cutoff to identify positive cells.

### Cell-type analyses

Cell-type analyses were conducted using data from a recently published dataset of a single-cell RNA sequencing atlas of healthy adult mouse liver [18].

### Statistical analyses

GraphPad Prism (version 9.20) was used for all statistical analyses. One-way ANOVA followed by Tukey’s multiple comparison tests were used to analyze differences between zonal expression.

## Results

### Establishing metabolic zones using multiplex RNAscope

We mapped the mRNA expression of key components of the Wnt signaling machinery in a zone-specific manner via multiplex RNAscope. First, we confirmed RNAscope’s capability to measure zonated expression by focusing on zonated markers, Gls2 and GS, in adult mouse liver. Expression of these markers was quantified in 20µm increments across a metabolic unit, beginning at the midline of the central vein (labelled 0µm) and ending at the midline of the portal vein (labelled 360µm) (Fig. S1). We observed a gradient of Gls2 mRNA expression progressively increasing towards the portal vein (Fig. S1A). By contrast, GS expression is most enriched around the central vein (Fig. S1B). We discovered that dividing metabolic units into thirds according to our 20µm increments relative to the central vein accurately defined the zonated expression of Gls2 and GS as Zone 1 and Zone 3 markers, respectively (*e*.*g*., 20-120µm, zone 1; 140-260 µm, zone 2; 280-360µm, zone 3) (Fig. 1A, 1B, Fig. S1C). These data are consistent with earlier immunohistochemical and functional studies of these enzymes [19, 20]. Together, our results demonstrate that multiplex RNAscope is a valid approach to measure zonated gene expression *in situ*.

**Fig. 1.**
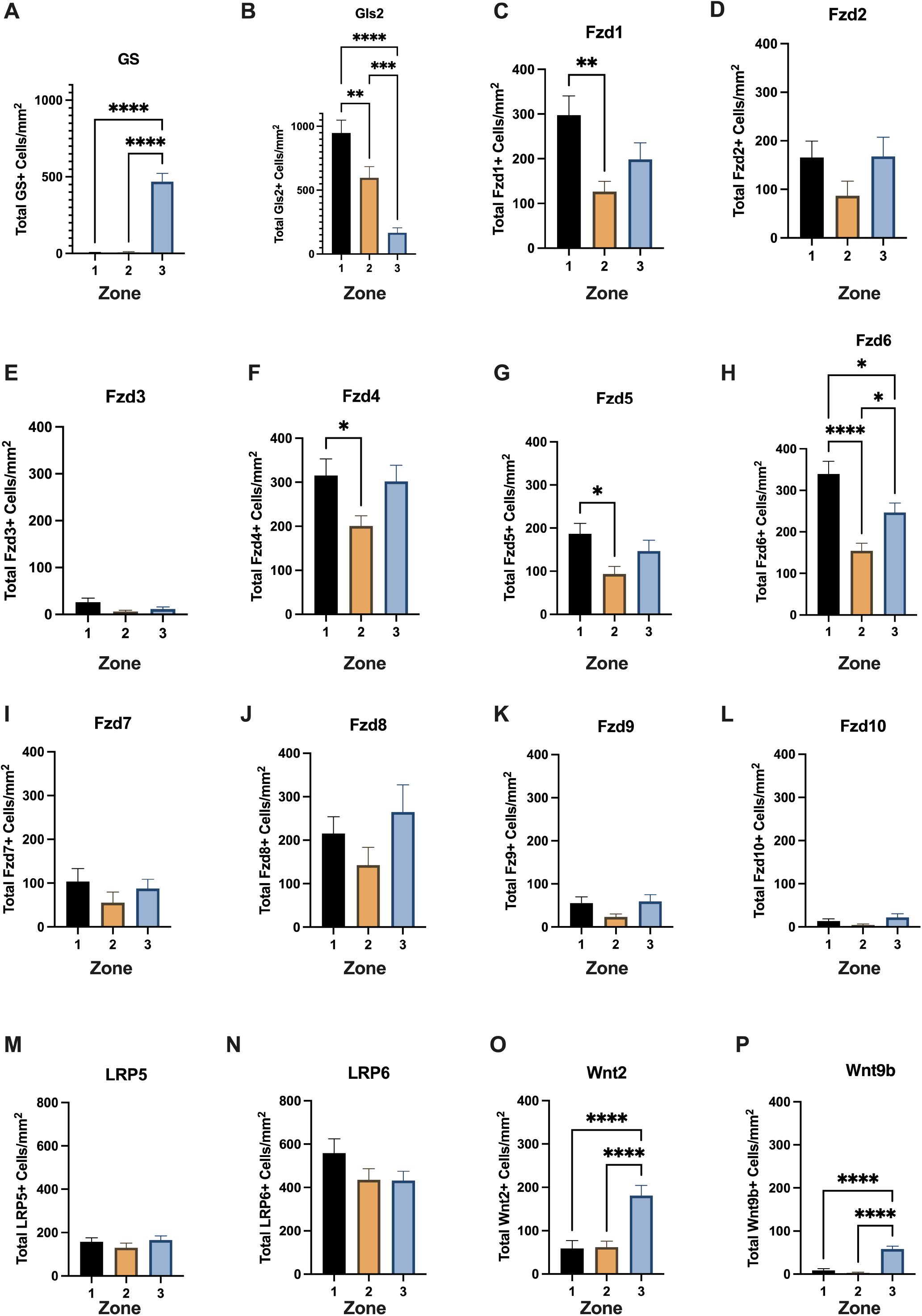
Hepatic expression of Wnt/ß-catenin signaling pathway components. (A, B) mRNA expression via multiplex RNAscope of (A) glutamine synthetase (GS) reveals strongly zonated expression in zone 3. One-way ANOVA with *post hoc* analyses. F(2, 79) = 70.64, p<0.0001. (B) Glutaminase2 (Gls2) is expressed along a gradient, significantly decreasing from zone 1 towards zone 3. One-way ANOVA with *post hoc* analyses. F(2, 77) = 25.10, p<0.0001)]. (C-L) mRNA expression of hepatic Frizzled receptors (Fzds) 1-10 via multiplex RNAscope. There is significantly higher zone 1 expression of: (C) Fzd1 [(F(2, 84) = 5.884, p=0.004)]; (F) Fzd4 [(F (2, 87) = 3.587, p=0.031)]; and (G) Fzd5 [(F (2, 84) = 4.234, p=0.017)] compared to zone 2. There is no significant zonation of: (D) Fzd2, (E) Fzd3, (I) Fzd7, (J) Fzd8, (K) Fzd9, and (L) Fzd10 (p>0.05). In contrast, (H) Fzd6 displays significant zonation, with highest expression in zone 1 [(F (2, 84) = 14.19, p<0.0001)]. (M, N) mRNA expression of Wnt ligands, (M) Wnt2 [(F (2, 84) = 13.54, p<0.0001)] and (N) Wnt9b [(F (2, 81) = 39.76, p<0.0001)] is strongly zonated with highest expression in zone 3. (O, P) mRNA expression of Fzd co-receptors low-density lipoprotein receptor-related protein 5/6 (lrp5/6). Neither (O) Lrp5 or (P) Lrp6 show zonated expression (p>0.05) Mean±SEM, n=3 mice for all conditions. **p<0.01, ***p<0.001, ****p<0.0001.

### Quantification of hepatic Frizzled receptor and Lrp5/6 co-receptor expression

While zonated Wnt signaling is established in adult liver, the precise identities of the cognate Fzd receptors responsible for this signaling have remained unknown. We therefore used RNAscope to determine Fzd expression at the mRNA level, and whether these receptors are zonated. We measured the number of cells positive for each respective Fzd receptor and found that all 10 Fzds are expressed in adult mouse liver, albeit with varying levels of expression (Fig.1C-L). Fzd1, Fzd4, and Fzd6 demonstrate highest levels of expression (Fig.1C, 1F, 1H), while Fzd3, Fzd9, and Fzd10 are expressed at low, but detectable levels (Fig. 1E, 1K, 1L). Moreover, Fzd2, Fzd5, Fzd7, and Fzd8 are expressed at levels intermediate to either extreme (Fig.1D, 1G, 1I, 1J). Surprisingly, all Fzds, with the exception of Fzd6, are not zonated. By contrast, Fzd6 is preferentially expressed in Zone 1 (Fig. 1H, Fig. S2). We also measured the average mRNA grains per cell for each respective Fzd (Fig. S3). mRNA grain number similarly reflected the same patterns as Fzd^+^ cell number (Fig. S3A-J), suggesting that levels of expression per cell are not directly influenced in a zone-specific manner.

We also analyzed mRNA expression of Fzd co-receptors Lrp5 and Lrp6. Though Lrp5 and Lrp6 are both expressed, neither display zonation. Interestingly, Lrp6 expression in all three zones is 3-fold higher compared to Lrp5 (Fig. 1M, 1N). Consistent with this, the levels of Lrp6 mRNA per cell are also ∼2-fold higher versus Lrp5 (Fig. S3K, S3L).

### Hepatic Wnt2 and Wnt9b expression is zonated

Since there is overwhelming evidence that metabolic zonation is driven by Wnt- and Fzd-dependent signaling, we hypothesized that the absence of zonated expression of most Fzds suggests that zonation is ligand-driven. Therefore, we posited that expression of Wnt ligands is zonated with preferential expression in zone 3, where the highest levels of Wnt signaling occur [21]. We specifically focused on Wnt2 and Wnt9b since they are critical for pericentral β-catenin activation and maintenance of zonation [12, 22]. We found that Wnt2 and Wnt9b mRNA expression is significantly elevated in zone 3 relative to the other zones, consistent with zonation of both Wnts (Fig. 1O, 1P). Additionally, Wnt2 expression is markedly higher versus Wnt9b, including in zone 3. Finally, numbers of mRNA grain per cell matched our above data (Fig. S3M, S3N). Together, these results support the importance of Wnt expression in driving hepatic zonation.

### Analysis of Wnt/Fzd signaling machinery by hepatic cell type

To determine the identities of hepatic cell types that express the machinery of Wnt/Fzd signaling, we analyzed single-cell RNAseq data of adult mouse liver [18]. We first analyzed cell-type specificity for expression of Fzds 1-10 and found each Fzd receptor possesses a distinct cell-type profile (Fig. 2A-J). Among the highest expressed Fzds (Fzd1, Fzd4, Fzd6), Fzd4 and Fzd6 are expressed in hepatocytes, alongside non-parenchymal cells including endothelial cells, cholangiocytes, and fibroblasts (Fig. 2D, 2F). In contrast, Fzd1 is not expressed in hepatocytes, but is robustly expressed in conventional type 1 dendritic cells (cDC1s) (Fig. 2A). Among the least expressed Fzds (Fzd3, Fzd9, Fzd10), Fzd3 and Fzd9 are mainly expressed in cholangiocytes, whereas negligible amounts of Fzd10 are found in stromal cells and plasmacytoid dendritic cells (pDCs) (Fig. 2C, 2I, 2J). Fzds with intermediate expression (Fzd2, Fzd5, Fzd7, Fzd8) display the broadest cell-type distribution with expression in hepatocytes, as well as a variety of non-parenchymal cells (Fig. 2B, 2E, 2G, 2H). While it was previously established that Wnt2 and Wnt9b are secreted by endothelial cells [12], we find that Wnt2 is not only expressed in endothelial cells, but in a variety of cell types as well, including hepatocytes and Kupffer cells (Fig. 2K). In contrast, Wnt9b is predominantly expressed in endothelial cells (Fig. 2L).

**Fig. 2.**
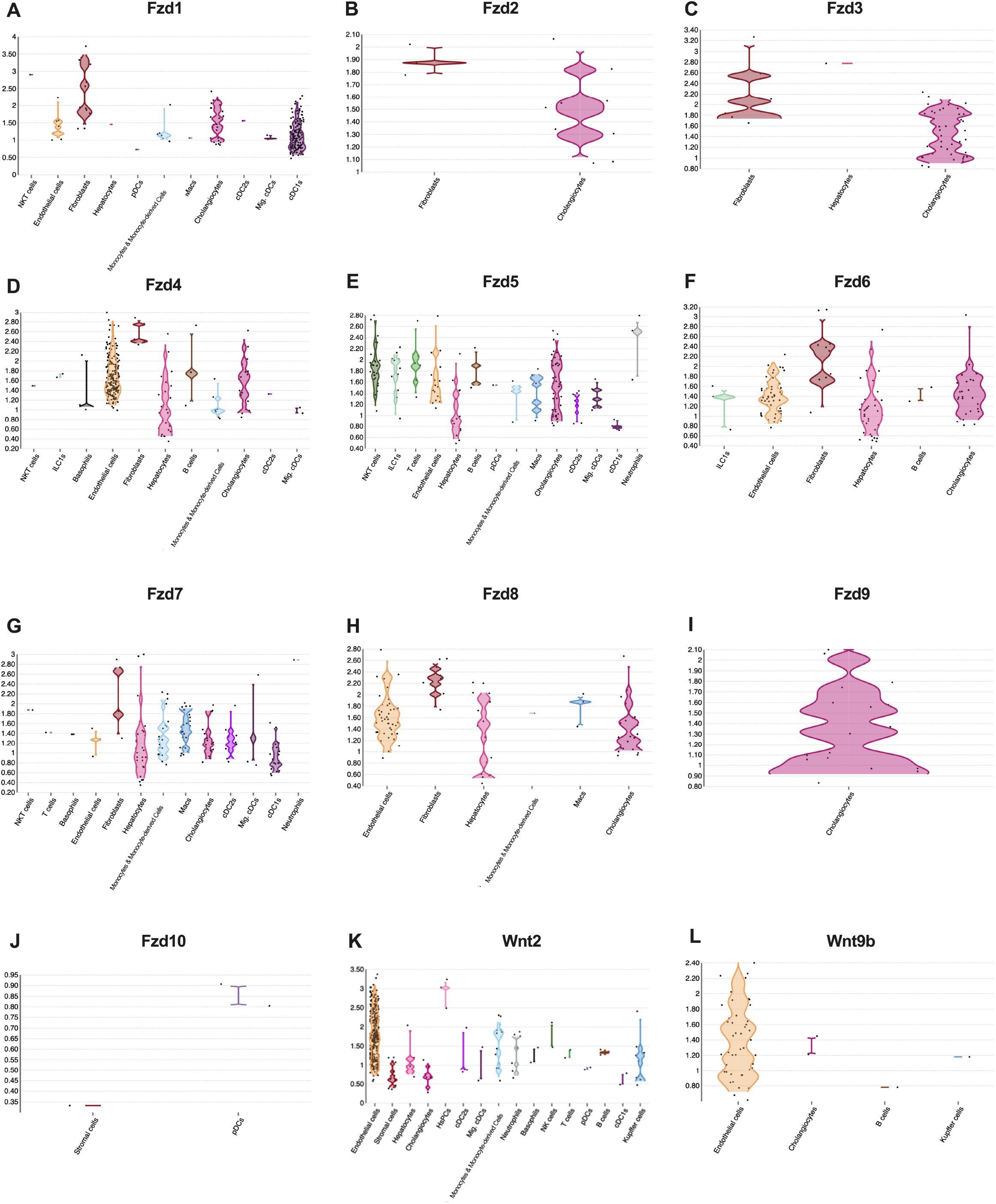
Cell-specific expression of Wnt signaling components. (A-H) Single-cell RNAseq revealsl Fzd receptor expression is present throughout a majority of different liver cell types, except for (I) Fzd9 which is solely expressed in cholangiocytes and (J) Fzd10 which is expressed in stromal cells and plasmacytoid dendritic cells (pDCs) only. (K) Wnt2 is expressed throughout various liver cell-types, whereas (L) Wnt9b is predominantly expressed in endothelial cells.

## Discussion

Wnt signaling is increasingly recognized as critical for regulation of hepatic zonation [23]. Until recently, studies of this signaling pathway at the protein level have been restricted due to a limited repertoire of antibodies capable of recognizing endogenously expressed Wnt/Fzd receptors. We overcame this limitation by establishing multiplex RNAscope in liver to define hepatic Wnt/Fzd expression. Our data indicates that all 10 Fzd receptors are expressed within a hepatic metabolic unit. Moreover, the Fzds are expressed in very different ways – in both the magnitude of expression and cell-type specificities.

We discovered that all Fzds, save for Fzd6, are expressed in a non-zonated manner. Interestingly, Fzd6 is preferentially expressed in Zone 1, the region exhibiting the least Wnt/beta-catenin signaling activity [21]. How does one reconcile this apparent discrepancy, *i*.*e*., expressing high levels of a Fzd in a region with the least Wnt signaling activity? Past work has shown that, unlike the other Fzds, Fzd6 does not transduce canonical Wnt signaling. Rather, Fzd6 functions as a negative regulator of Wnt signaling as it lacks a C-terminal PDZ-binding domain responsible for transducing Wnt-dependent downstream signaling [24, 25]. We therefore postulate that Fzd6’s repressive properties enhance zonation by further restricting Wnt signaling to Zone 3. Conversely, it is possible that the expression of the other Fzds becomes increasingly zonated post-development in response to hepatic injury to facilitate cellular regeneration. Future work is clearly required to explore such possibilities.

In contrast to the non-zonated expression patterns of most Fzd receptors, Fzd ligands, Wnt2 and Wnt9b are both highly zonated. This is consistent with earlier work showing that Wnt/beta-catenin signaling is most evident in zone 3 [12]. This suggests that maintenance of hepatic zonation is driven by ligand availability rather than the expression of the receptors themselves under physiological conditions.

Our results also show that cell-type specific expression of Fzds1-10 as well as Wnt2 and Wnt9b is not solely restricted to hepatocytes, but is found in a variety of non-parenchymal cells as well. Previously, we showed that endothelial cells predominantly secrete Wnt2 and Wnt9b [12]. In fact, we eliminated expression of Wnt2 and Wnt9b from endothelial cells of the liver and identified the observed phenotype to mimic liver-specific-β- catenin knockouts (KO), liver-specific LRP5-6-double KO and endothelial-specific Wntless KO in terms of loss of zone 3 gene expression and delay in liver regeneration after partial hepatectomy [12, 13, 26, 27]. However, here we found that there are significant differences in the cell-type specific expression of these Wnts. While Wnt9b expression is primarily restricted to endothelial cells, Wnt2 is more broadly expressed across several hepatic cell types (*e*.*g*., hepatocytes and monocytes). This suggests that these Wnts may play distinct roles in maintenance of zonation based on the spatial distribution of these cell types. Cell-type specific genetic manipulation could further elucidate the respective contributions of these Wnts. Furthermore, we also find that many of the Fzds are robustly expressed in endothelial cells, suggesting significant autocrine Wnt/Fzd signaling locally within the hepatic vasculature. Indeed, local endothelial Wnt signaling maintains endothelial cell identity and zonation [28]. Moreover, these data are consistent with earlier work in brain demonstrating a role for Wnt/β-catenin signaling in vascular development and maintenance post-development [29].

Limitations include the possibility that our mRNA expression findings do not directly translate to the protein level. Potential differences in mRNA stability or post-translational modifications may alter expression of Fzd receptor proteins. On the other hand, several molecules we assayed by RNAscope (*e*.*g*., GS, Gls2, Wnt2, Wnt9b) have matched the spatial distribution at the protein level [12, 30]. Additionally, our study focused on male mice, making future work in females critical for identifying possible sex differences.

In summary, the results presented here provide the first comprehensive map of Fzd receptor mRNA expression in adult liver, alongside additional components of the Wnt signaling pathway. We find that all 10 Fzds are expressed within metabolic units, alongside Lrp5 and Lrp6. However, only Fzd6 is expressed in a zone-specific manner. Furthermore, we show that Wnt2 and Wnt9b are strongly zonated which points to the importance of ligand availability in driving zonated expression.

## Acknowledgements

This work was supported by the National Institutes of Health (R01DK062277-17S1, T32GM133353 to J.G., and P30DK120531, R21052419, R21AA028800, AG068607, R01DK124219 to Z.F.) and the Department of Defense (PR141292, PR210207 to Z.F.). This work was also supported in part by 1R01DK62277 and 1R01DK100287 to S.P.M.

## Author contributions

The study was designed and conceptualized by Z.F., S.P. M., and J.G. Experiments were conducted by J.G., S.H., E.D., and P.J. Data interpretation were conducted by J.G., S.L., and Z.F. The manuscript was written by J.G., Z.F, and S.P. M., with input from the other authors.

