## Supplementary Information for "A spatial atlas of Wnt and Frizzled receptor expression in adult mouse liver"

**Gayden *et al.***

**Supplementary Figures S1-S3**

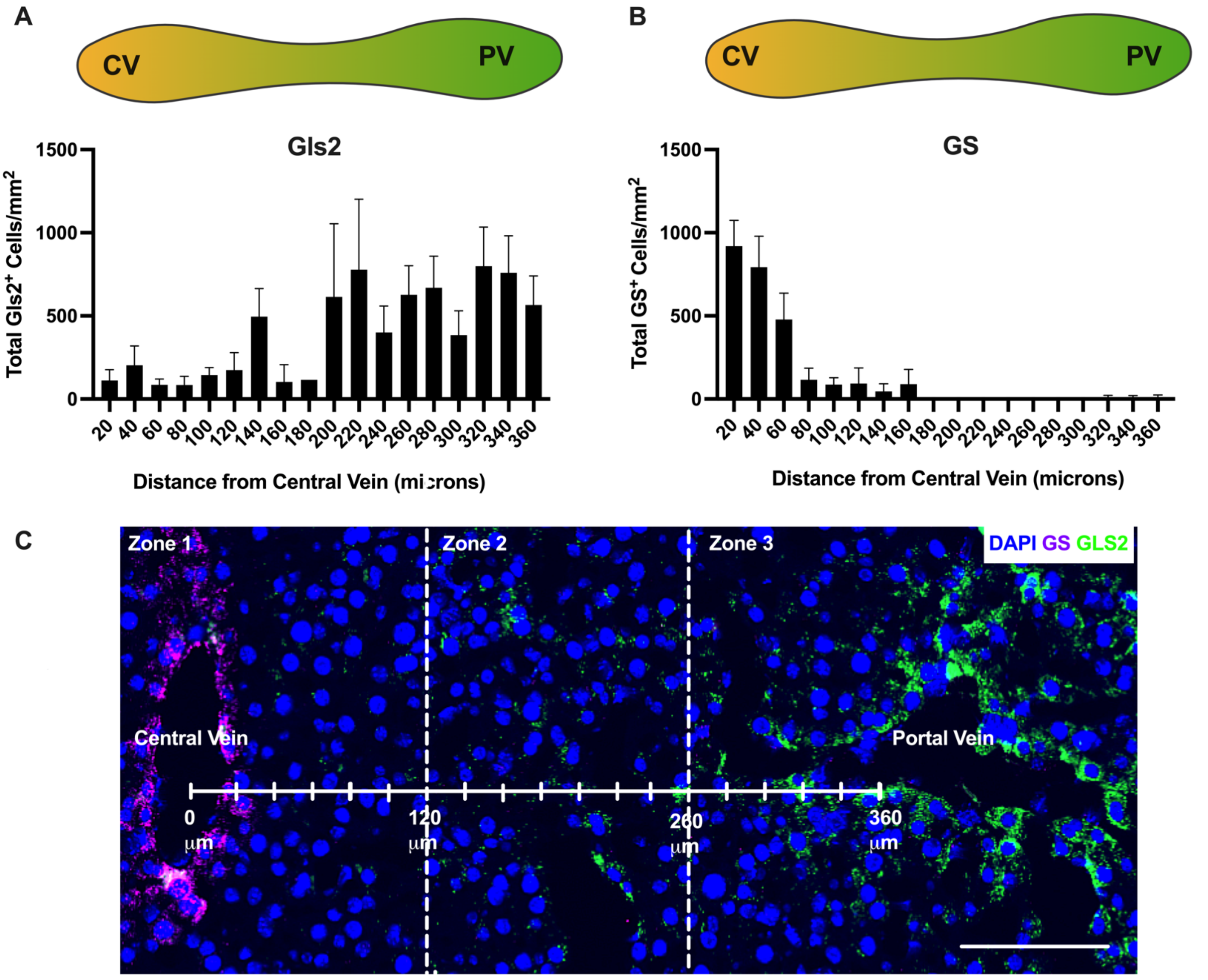


**Fig. S1. Establishment of metabolic zones using multiplex RNAscope.** (A, B) Quantification of cells positive for glutaminase2 (Gls2) and glutamine synthetase (GS) mRNA expression within a metabolic unit is quantified in 20µm increments, starting at 20µm from the midline of the central vein (CV, labelled 20µm) and terminating at the midline of the portal vein (PV, labelled 360µm). (A) Gls2 expression progressively increases towards the periportal region, whereas (B) GS expression is restricted to the pericentral region. (C) Representative metabolic unit with GS (purple) and Gls2 (green) outlining the CV and PV, respectively. Cell nuclei are labeled with DAPI stain (in blue). Scale bar = 100µm. Mean±SEM, n=3 mice for all conditions.

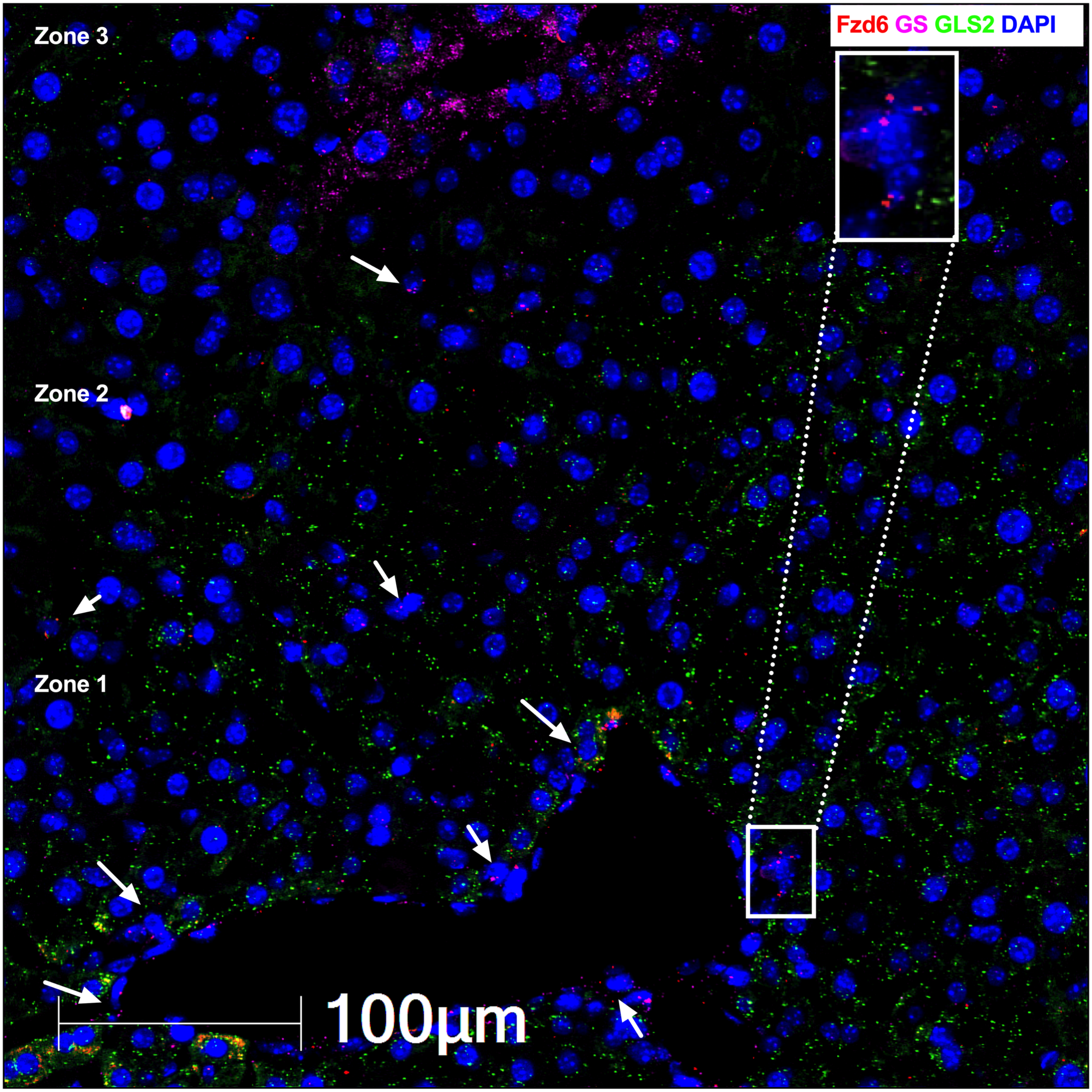


**Fig. S2. Representative image of Fzd6 mRNA expression in adult mouse liver.** Confocal image of Fzd6 mRNA alongside GS and Gls2 marking the central vein and portal veins, respectively. Imaging reveals Fzd6 is most expressed in hepatic zone 1. Cells expressing Fzd6 are indicated by arrows. A zoomed-in Fzd6$^{+}$ cell is featured within the inset. Scale bar=100µm.

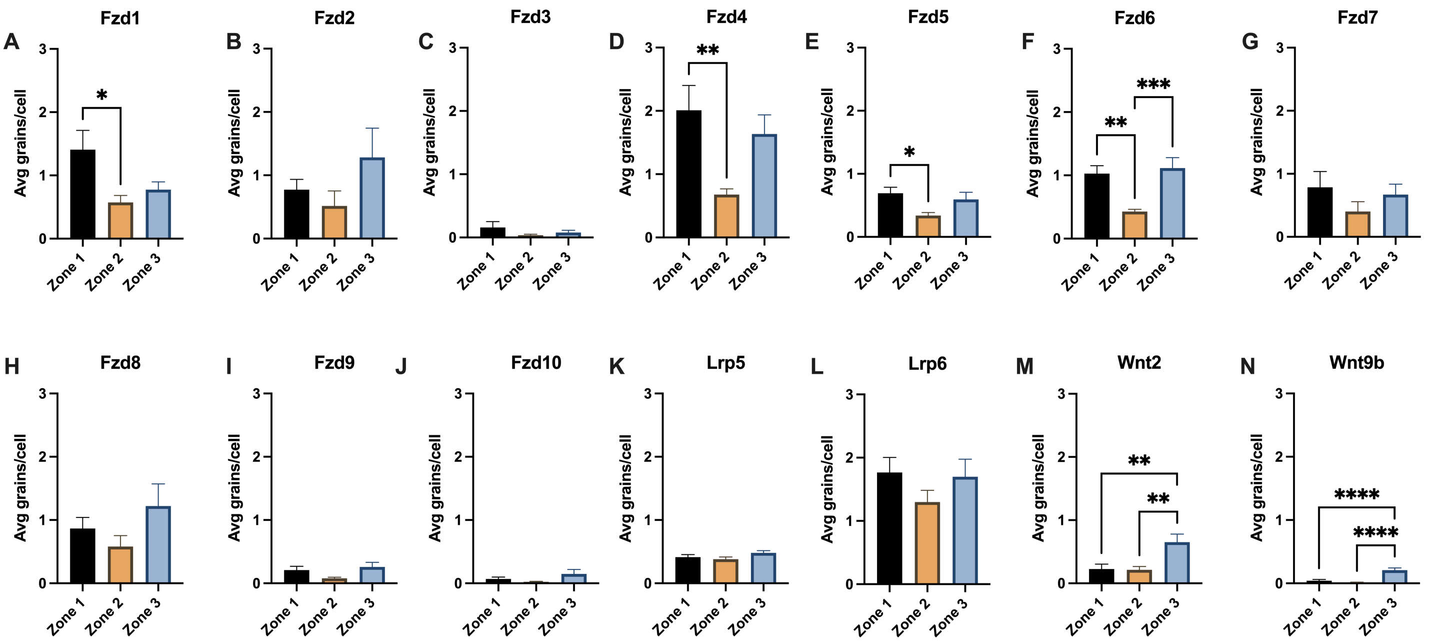


**Fig. S3. Quantification of mRNA grains per cell.** Quantitative analysis of mRNA grain number corresponding to (A-H) Frizzled receptors 1-10, (K, L) Wnt2, Wnt9b, and (M, N) Lrp5 and Lrp6. *$p<0.01$, **$p<0.001$, ***$p<0.0001$, ****$p<0.00001$. Mean±SEM, n=3 mice for all conditions.
